# Proteome reallocation enables the selective *de novo* biosynthesis of non-linear, branched-chain acetate esters

**DOI:** 10.1101/2022.03.17.484697

**Authors:** Hyeongmin Seo, Richard J. Giannone, Yung-Hun Yang, Cong T. Trinh

## Abstract

The one-carbon recursive ketoacid elongation pathway is responsible for making various branched-chain amino acids, aldehydes, alcohols, and acetate esters in living cells. Controlling selective microbial biosynthesis of these target molecules at high efficiency is challenging due to enzyme promiscuity, regulation, and metabolic burden. In this study, we present a systematic modular design approach to control proteome reallocation for selective microbial biosynthesis of branched-chain acetate esters. Through pathway modularization, we partitioned the branched-chain ester pathways into four submodules including keto-isovalerate submodule for converting pyruvate to keto-isovalerate, ketoacid elongation submodule for producing longer carbon-chain keto-acids, ketoacid decarboxylase submodule for converting ketoacids to alcohols, and alcohol acyltransferase submodule for producing branched-chain acetate esters by condensing alcohols and acetyl-CoA. By systematic manipulation of pathway gene replication and transcription, enzyme specificity of the first committed steps of these submodules, and downstream competing pathways, we demonstrated selective microbial production of isoamyl acetate over isobutyl acetate. We found that the optimized isoamyl acetate pathway globally redistributed the amino acid fractions in the proteomes and required up to 23-31% proteome reallocation at the expense of other cellular resources, such as those required to generate precursor metabolites and energy for growth and amino acid biosynthesis. The engineered strains produced isoamyl acetate at a titer of 8.8 g/L (> 0.25 g/L toxicity limit), a yield of 0.17 g/g (47% of maximal theoretical value), and 86% selectivity, achieving the highest titers, yields and selectivity of isoamyl acetate reported to date.

## INTRODUCTION

Short-chain esters formulate volatile compounds commonly found in flowers, ripe fruits, and fermenting yeasts (Sugimoto et al., 2021; Sumby et al., 2010). Some of these esters are suggested to have an important ecological role in pollination (Knudsen and Tollsten, 1993). Industrially, these esters have versatile utility as flavors, fragrances, solvents, and biofuels. For instance, isoamyl acetate (3-methyl-1-butyl acetate) is known as banana oil with a global market value of $5 billion in 2019 (IndustryARC, 2019; IndustryResearch, 2021). An isomer of isoamyl acetate, ethyl valerate, is fully compatible for blending with gasoline or diesel (Lange et al., 2010), suggesting potential application of isoamyl acetate as drop-in biofuel. Currently, knowledge about the microbial biosynthesis of these molecules from renewable and sustainable feedstocks is limited, making it difficult to optimize their production without further interrogation.

Biologically, cells can synthesize an ester by condensing an alcohol and an acyl-CoA with an alcohol acyltransferase (AAT) (Mason and Dufour, 2000). Due to the abundance and essentiality of acetyl-CoA in living cells, acetate esters are the most common esters found in nature. By activating one-, two-, or three-carbon recursive elongation via the recursive fatty acid biosynthesis (Liu et al., 2016; Youngquist et al., 2013), reverse beta-oxidation (Dellomonaco et al., 2011), or Ehrlich pathways (Atsumi et al., 2008; Zhang et al., 2008), it is possible to synthesize a large library of acetate esters containing unique alcohol moieties with linear, branched, even, and/or odd carbon chains (Layton and Trinh, 2016a; Layton and Trinh, 2016b; Lee and Trinh, 2020). However, selective microbial biosynthesis of designer acetate esters at high efficiency has been an outstanding metabolic engineering problem. For instance, branched-chain acetate esters (e.g., isoamyl acetate) represent an important class of molecules that can be synthesized via the one-carbon recursive ketoacid elongation pathway (Connor et al., 2010; Connor and Liao, 2008).

Starting from the precursor pyruvate, this pathway generates ketoacids that can be decarboxylated to aldehydes, reduced to branched-chain alcohols, and condensed to acetate esters. Isobutyl acetate is generated in the first cycle, followed by isoamyl acetate in the second cycle, and so on. Although microbial production of isoamyl acetate has been reported since early 2000s by the condensation of isoamyl alcohol and acetyl CoA, production titers (<1 g/L) and selectivities (< 30%) were relatively low (Abe and Horikoshi, 2005; Horton et al., 2003; Vadali et al., 2004a; Vadali et al., 2004b).

Many confounding factors might negatively affect selective microbial biosynthesis of branched-chain acetate esters. In addition to the well-known toxicity of higher alcohols and esters (Wilbanks and Trinh, 2017b) and required expression of multiple pathway enzymes (Tai et al., 2015), the recursive one-carbon elongation pathway generates intermediate alcohol byproducts (e.g., isobutanol) that compete with the target biosynthesis of esters (e.g., isobutyl acetate instead of isoamyl acetate) (Dellomonaco et al., 2011; Layton and Trinh, 2014; Marcheschi et al., 2012; Martin et al., 2003; Zhang et al., 2008). Currently, understanding and controlling this recursive elongation pathway for efficient biosynthesis of target branched-chain acetate esters remains elusive. A cellular proteome constitutes ∼50% of dry cell weight, requiring a significant resource investment (Neidhardt et al., 1990). As rewiring cellular metabolism can severely impact overall proteome allocation, especially when multiple enzyme pathways are introduced and/or overexpressed, proteome allocation or reallocation must be considered to achieve optimal product production. This reallocation, however, is complex and poorly understood because it requires a precise control of the expression, specificities, and activities of multiple pathway enzymes in order to achieve optimal metabolic fluxes for selective microbial production of the target molecule (Lechner et al., 2016) and avoid metabolic burden (Wu et al., 2016).

In this study, we presented a systematic modular design approach to control proteome reallocation for the selective microbial biosynthesis of branched-chain acetate esters via manipulation of substrate specificity and expression level of multiple pathway enzymes. For proof-of-concept, we demonstrated the approach to enable selective production of isoamyl acetate over isobutyl acetate by controlling the one-carbon recursive ketoacid elongation pathway. Using quantitative proteomics, we shed light on pathway-level proteome reallocation, metabolic burden, and bottlenecks, which guided the effective metabolic rewiring for the efficient target ester biosynthesis.

## RESULTS

### Design, construction, and characterization of a generalizable modular, non-linear, branched-chain acetate ester pathway

#### Modular pathway design principles

The branched-chain acetate ester biosynthesis pathway is derived from pyruvate (Fig. 1a). Pyruvate is converted to 2-ketoisovalerate via the L-valine biosynthesis pathway (KIV submodule) then elongated to 2-ketoacids via the +1 recursive ketoacid elongation cycle mediated by the LeuABCD operon (keto-acid elongation submodule). The Ehrlich pathway (KDC submodule) converts 2-ketoacids to aldehydes and alcohols, then the alcohol acyltransferase pathway (AAT submodule) condenses alcohols and acetyl-CoA to form branched-chain acetate esters. The key enzymes governing each pathway submodule such as acetolactate synthase (IlvB or AlsS) (Steinmetz et al., 2010), 2-isopropylmalate synthase (LeuA) (Ulm et al., 1972; Wiegel and Schlegel, 1977), ketoacid decarboxylase (KDC) (de la Plaza et al., 2004; Mak et al., 2015), and alcohol acyltransferase (AAT) (Cumplido-Laso et al., 2012; Nancolas et al., 2017) are promiscuous, formulating a generalizable non-linear, branched-chain acetate ester biosynthesis pathway derived from the interconnected submodules (Fig. 1b). Due to the enzyme promiscuity and pathway modularity, we hypothesized that balancing proteome of the individual submodules with manipulated key enzymes (i.e., AlsS, LeuA, Kdc, and AAT) is critical to control selectivity of designer acetate ester production with high titers and yields. As a proof of concept, we demonstrated the feasibility of controlling selective production of isoamyl acetate over isobutyl acetate as a byproduct.

**Figure 1.**
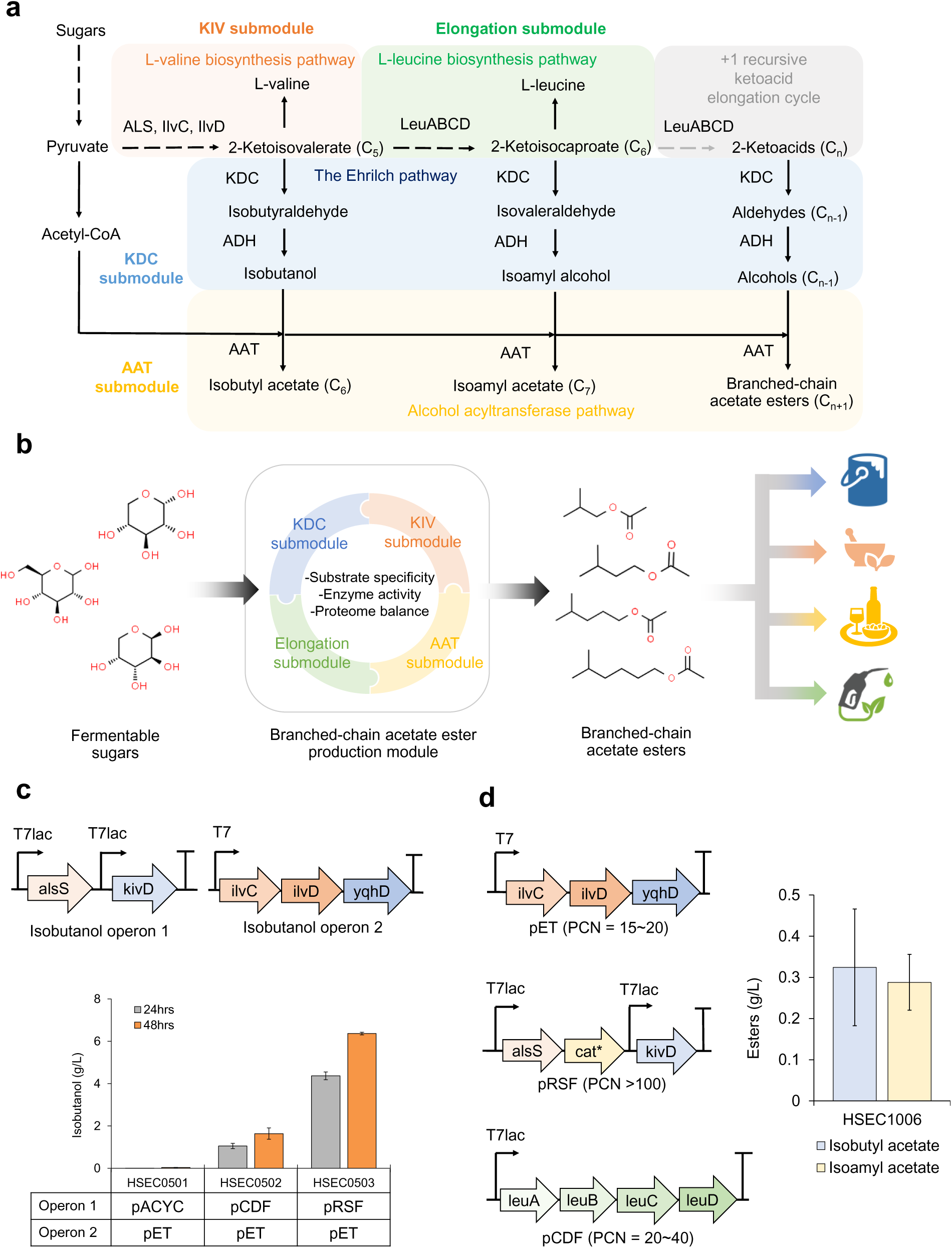
Modular design of branched-chain acetate ester production pathway. **(a)** A metabolic map of branched-chain acetate ester biosynthesis pathway. **(b)** The four submodules of the branched-chain acetate ester pathway designed for controlling selective microbial biosynthesis of acetate esters useful for various industries. **(c)** Optimization of isobutanol production module. Shown are the genetic architecture of the two isobutanol operons and characterization of isobutanol production by varying plasmid copy numbers (PCNs) in the engineered strains. Plasmid copy numbers of pRSF, pCDF, pET, and pACYC are >100, 20∼40, 15∼20, and 10∼12, respectively. **(d)** Genetic architecture of operons designed for isoamyl acetate production and design validation. Each data in panels **(c)** and **(d)** represents a mean ± 1 standard deviation from at least three biological replicates.

#### Construction and characterization of KIV and KDC submodules

We first aimed to construct a strong KIV submodule to produce 2-ketoisovalerate (KIV), the key precursor metabolite of the branched-chain acetate ester pathway. Because metabolic balance between the KIV and KDC submodules is also important for high production of branched-chain alcohols, we started by optimizing the expression of these two submodules simultaneously that can be evaluated by measuring isobutanol production. Since heterologous expression of acetolactate synthase (AlsS) from *Bacillus subtilis* and 2-ketoisovalerate decarboxylase (KivD) from *Lactococcus lactis* is critical for high-level isobutanol production (Atsumi et al., 2008), we constructed three different plasmids (i.e., pACYC, pCDF, and pRSF) harboring *alsS* and *kivD* with different plasmid copy numbers (PCNs) to rapidly optimize the production submodules. Then, we introduced them to a BL21 (DE3) strain harboring an operon of *ilvC*, *ilvD*, and *yqhD* under the control of T7 promoter in a medium copy number pET23a plasmid (Fig. 1c). Isobutanol production by the recombinant *E. coli* strains were measured to identify the most efficient combination of the production submodules. The result showed that HSEC0503 harboring *alsS* and *kivD* in a high copy number plasmid (pRSFDuet-1) produced the highest level of isobutanol (6.4 g/L) within 48 hours (h), while HSEC0501 carrying *alsS* and *kivD* in a low copy number plasmid (pACYCDuet-1) produced only 0.03 g/L of isobutanol (Fig. 1c). This result suggested that the high-level expression of *alsS* and *kivD* to increase enzyme levels is important to pull the metabolic flux towards the branched-chain acetate ester pathway.

#### Assembly and characterization of the de novo isoamyl acetate pathway module

Combining the elongation and AAT submodules together with the KIV and KDC submodules forms the isoamyl acetate pathway (Figs. 1a, 1b, 1d). To construct the elongation submodule, a *leuABCD* operon was cloned in a medium copy number pCDFDuet-1 plasmid (Fig. 1d). While overexpressing the elongation submodule pulls metabolic flux towards isoamyl acetate, it should be noted that isobutyl acetate can still be produced as a byproduct due to the promiscuity of the KDC submodule that can produce either isobutanol or isoamyl alcohol (de la Plaza et al., 2004). To enhance selective production of isoamyl acetate in our design, we reasoned that high substrate specificity towards isoamyl alcohol should be chosen to create the AAT submodule. Recently, a chloramphenicol acetyltransferase (CAT) was engineered and repurposed as an AAT to produce a broad range of esters, capable of converting isoamyl alcohol to isoamyl acetate with 95% (mol/mol) efficiency in *E. coli* (Seo et al., 2021). Specifically, the engineered CATec3 Y20F derived from *E. coli* exhibited 3.5-folds higher catalytic efficiency (k_cat_/K_M_) towards isoamyl alcohol than isobutanol (Seo et al., 2021), showing a higher selectivity against isoamyl alcohol than isobutanol as compared to other available AATs (Tai et al., 2015) (Fig. S1). Therefore, we deployed CATec3 Y20F to build the AAT submodule for selective production of isoamyl acetate (Fig. 1d).

By introducing the KIV, elongation, KDC, and AAT submodules into *E. coli* BL21(DE3), we created the strain HSEC1006 capable of performing the *de novo* biosynthesis of isoamyl acetate from fermentable sugars. The characterization results showed that HSEC1006 produced 0.08 g/L ethyl acetate, 0.32 g/L isobutyl acetate and 0.29 g/L isoamyl acetate from glucose with 44.3% selectivity (Fig. 1d). While the *de novo* isoamyl acetate biosynthesis was successfully demonstrated, the titer and selectivity were still low and hence required further pathway optimization.

### Enhancing selective production of isoamyl acetate module

To boost isoamyl acetate production with higher selectivity, we applied three push-and-pull metabolic engineering strategies: deletion of competing pyruvate and acetyl-CoA pool related genes (i.e., *adhE* encoding aldehyde-alcohol dehydrogenase used for ethanol synthesis and *dld* encoding quinone-dependent D-lactate dehydrogenase used for lactate synthesis), overexpression of a feedback insensitive LeuA G462D mutant (Mikhail Markovich Gusyatiner, 1999), and overexpression of a longer chain keto-acid specific KivD F381L/V461A (Zhang et al., 2008). We constructed four strains by implementing combinations of these strategies and measured acetate ester production (Fig. 2a). Deletion of *adhE* and *dld* in the strain HSEC1207 improved isoamyl acetate by 1.9-fold (0.5 g/L) as compared to HSEC1006, while isobutyl acetate titer was not significantly changed regardless of the increased isobutanol (Fig. 2c), suggesting that CATec3 Y20F was effective for selective isoamyl acetate production. A combination of KivD F381L/V461A overexpression and *adhE* and *dld* deletion in the strain HSEC1208 further improved isoamyl acetate titer up to 0.8 g/L, while isobutyl acetate production decreased to 0.2 g/L (Fig. 2a). Interestingly, HSEC1209 expressing feedback insensitive LeuA (G462D) and wild-type KivD produced the similar titers of isoamyl acetate as compared to HSEC1208, while only 0.06 g/L of isobutyl acetate was produced (Fig. 2a). The results indicate that both Kdc and LeuA enzymatic steps are critical for selective branched-chain acetate ester production.

**Figure 2.**
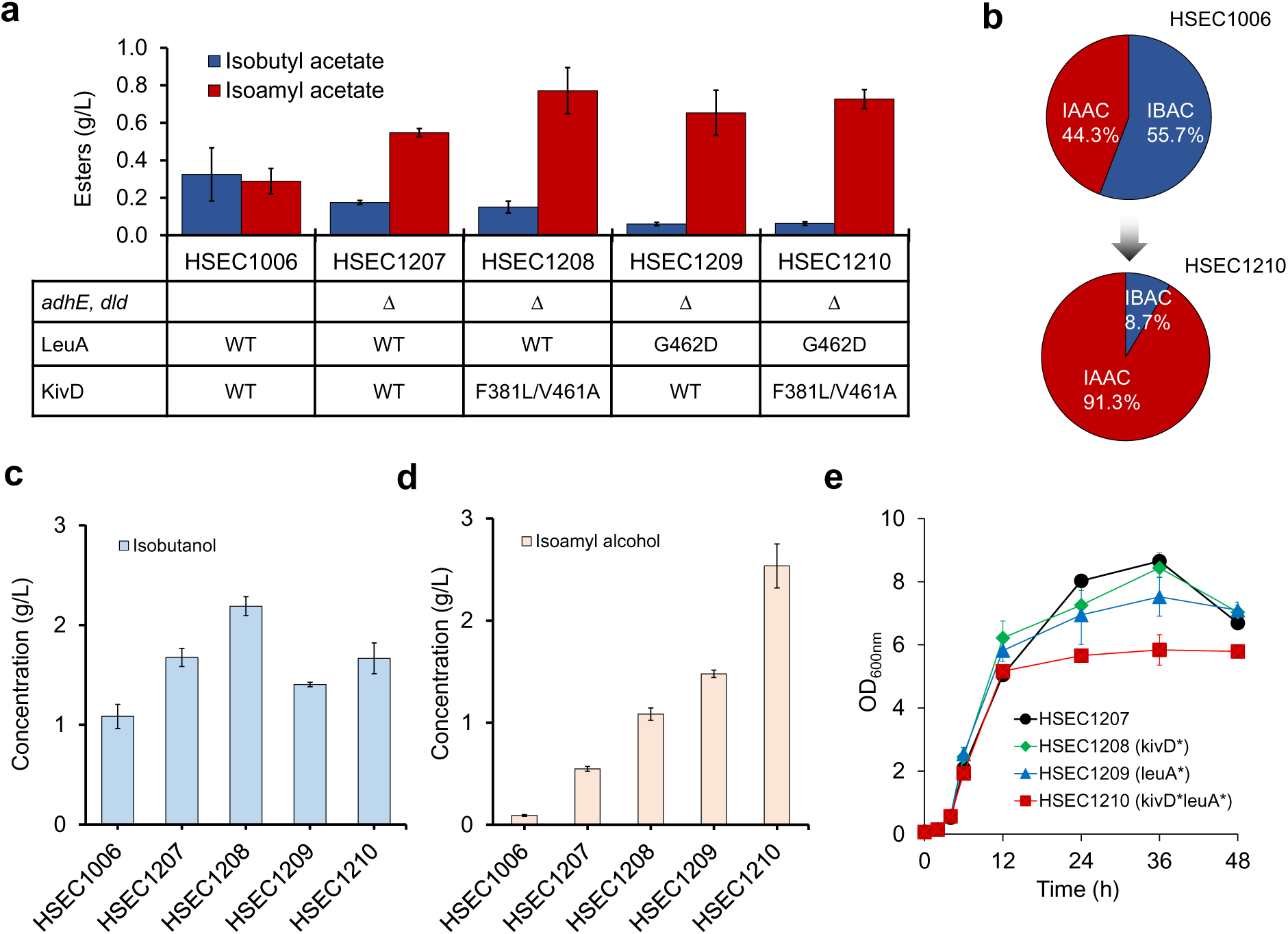
Controlling selective microbial production of isoamyl acetate. **(a)** Systematic characterization and comparison of push-and-pull metabolic engineering strategies for selective microbial biosynthesis of isoamyl acetate over isobutyl acetate. **(b)** Selectivity of isoamyl acetate production by HSEC1006 and HSEC1210. **(c-d)** Accumulation of isobutanol and isoamyl alcohol by the engineered strains. **€** Comparison of cell growth of the engineered strains during 48 h culturing. Each data represents mean ± 1 standard deviation from at least three biological replicates.

HSEC1210 expressing both KivD F381L/V461A and LeuA G462D did not improve isoamyl acetate production (Fig. 2a). Like HSEC1209, HSEC1210 significantly reduced the byproduct isobutyl acetate from 0.2 g/L to 0.07 g/L. Due to the reduced production of isobutyl acetate, selectivity of isoamyl acetate in the strain HSEC1210 increased 2.1-fold (91.3%), as compared to HSEC1006 (44.3%) (Fig. 2b). We observed that more than 92% (w/w) of isobutyl acetate and isoamyl acetate were extracted to the hexadecane layer, consistent with a previously reported extraction efficiency (Rodriguez et al., 2014). However, isobutanol and isoamyl alcohol were accumulated at a high level in HSEC1210, up to 1.5 g/L and 2.5 g/L in the culture medium, respectively (Figs. 2c, 2d). Because isoamyl alcohol at 2.5-3.0 g/L concentration inhibits 50-80% of cell viability (Connor and Liao, 2008; Wilbanks and Trinh, 2017a), the accumulated alcohols might have likely caused the 30% lower cell mass of HSEC1210 than the other strains (Fig. 2e).

Taken together, manipulating the key metabolic enzymes AlsS, Kivd, and LeuA can effectively control selective production of isoamyl acetate over isobutyl acetate. However, metabolic bottleneck(s) in the engineered pathway module is still present in HSEC1210, likely limiting its isoamyl acetate production.

### Proteomic analysis reveals proteome reallocation by isoamyl acetate pathway overexpression

#### Identification of metabolic bottlenecks due to imbalanced pathway protein allocation

Higher accumulation of alcohols in HSEC1210 implies that the metabolic flux of isoamyl acetate module was not well balanced. We hypothesized that this imbalance might have been caused by a metabolic burden imparted by the necessary overexpression of multiple heterologous and endogenous genes in the isoamyl acetate pathway. To understand this imbalance and identify the potential metabolic bottlenecks that might have limited isoamyl acetate production, we examined proteome reallocation of HSEC1210 growing under conditions with and without IPTG induction of the target pathway. As expected, the uninduced HSEC1210 (1.09 ± 0.03 1/h) grew faster than IPTG-induced HSEC1210 (0.63 ± 0.01 1/h) (Fig. 3a). Without the IPTG induction, HSEC1210 did not produce any detectable amount of isoamyl alcohol and isoamyl acetate likely due to the tight regulation of the isoamyl acetate pathway genes under the T7 and T7lac promoters. With IPTG induction, the isoamyl acetate biosynthetic proteins were significantly more abundant than the control, confirming all ten genes were successfully overexpressed (Figs. 3b, 3c). Especially, the three proteins AlsS, KivD*, and LeuA* represented an outsized share of protein expression relative to the whole proteome, exhibiting the highest levels of protein abundances in the pathway and were ranked at 1, 11, and 19 in increased abundances in the proteome, achieving 602-, 60-, and 47-fold more abundant than the control (without IPTG), respectively. These proteins represent three important metabolic steps to direct metabolic fluxes towards the high production of isoamyl acetate.

**Figure 3.**
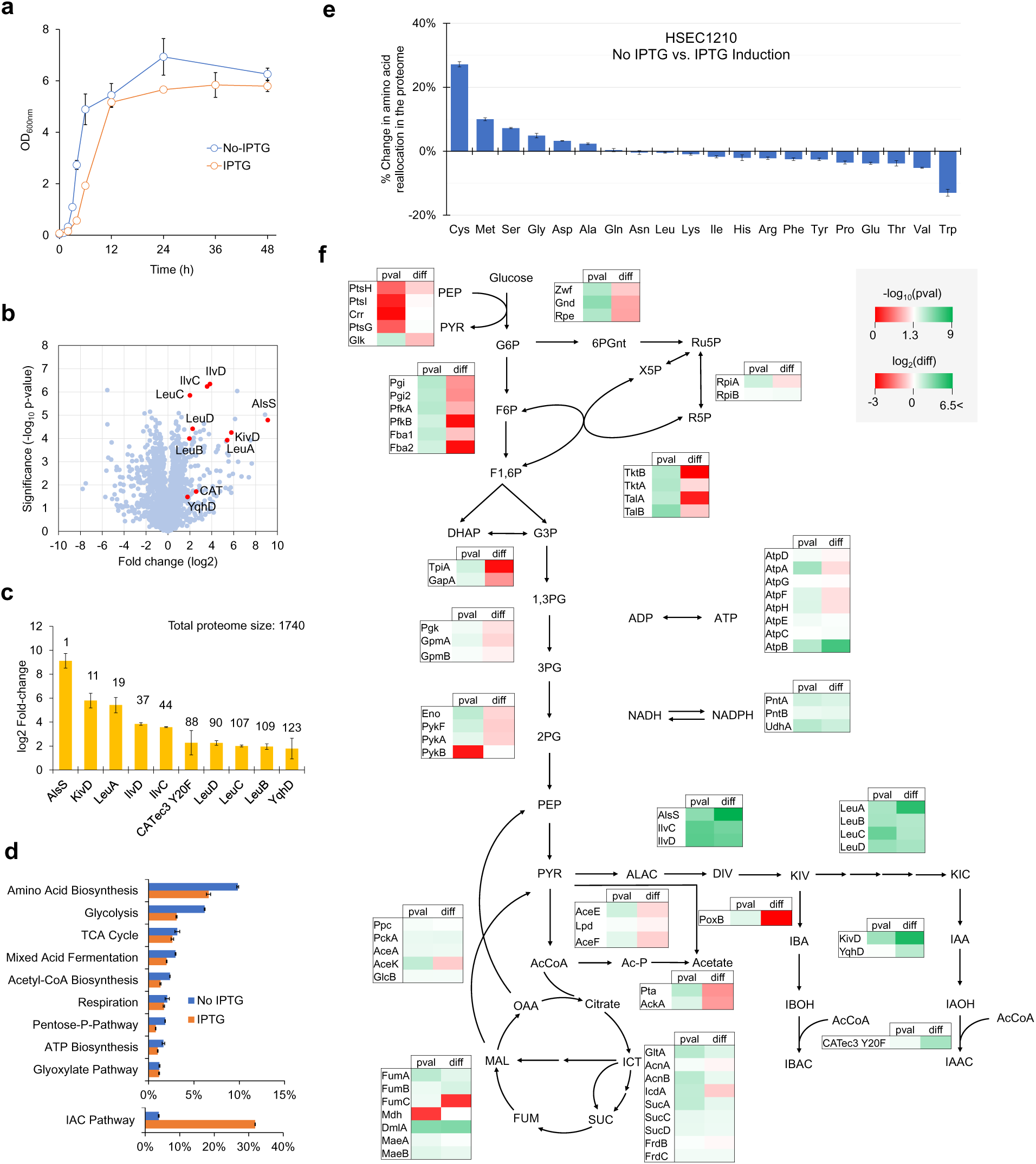
Analysis of proteome reallocation of HSEC1210 growing in media with and without IPTG induction. **(a)** Cell growth. **(b)** A volcano plot comparing HSEC1210 proteomes at 24h. The red dots indicate the isoamyl acetate pathway proteins. **(c)** Proteomic fold-change of enzymes in the isoamyl acetate pathway upon IPTG induction. (**d**) Mass fractions of the fueling pathways for generating precursor metabolites and energy and biosynthesis pathways of amino acid€**(e)** Percent of change in amino acid reallocation in the proteome upon IPTG induction. **(f)** Metabolic map displaying protein fold-changes of enzymes in the central metabolism. Significance (pval) and abundance fold-change (diff) were presented by –log_10_(p-value) and log_2_(difference), respectively. Each data represents mean ± 1 standard deviation from at least three biological replicates.

#### Overexpression of the isoamyl acetate module resulted in significant proteome reallocation

Upon IPTG induction, the mass fraction of the isoamyl acetate pathway proteome increased about 8-fold, representing 3.9% of total protein abundance in the uninduced control and 31.9% upon induction. This increase in protein abundances is seemingly at the expense of other cellular systems, pathways, and resources such as those invested for generation of precursor metabolites and energy (i.e., glycolysis, TCA cycle, mixed acid fermentation, acetyl CoA biosynthesis, respiration, pentose-P-phosphate, ATP biosynthesis, and glyoxylate pathway) and amino acid biosynthesis (Fig. 3d). For instance, by examining the glycolysis, we observed that the abundances of many glycolysis-related proteins, such as glucose-6-phosphate isomerase (Pgi), 6-phosphofructokinase (Pfk), fructose-and bisphosphate aldolases (Fba), were significantly reduced upon the isoamyl acetate pathway overexpression (Fig. 3f). The decreased abundances of the pyruvate dehydrogenase enzyme complex (i.e., AceE, Lpd, and AceF) of the acetyl-CoA biosynthesis pathway correlated well with the increased flux towards the isoamyl acetate pathway. Since 0.1mM IPTG concentration has little inhibitory effect on *E. coli* growth (Kosinski et al., 1992) and isoamyl alcohol accumulation (0.13 g/L) was relatively low during the exponential growth phase, the reduced growth was likely attributed to the metabolic burden caused by simultaneous overexpression of multiple enzymes.

By examining the isoamyl acetate pathway proteins, we could further identify an imbalanced overexpression among the target genes. Even though the target pathway genes that were expressed on higher copy plasmids yielded more abundant proteins as expected, the genes belonging to the same operons exhibited different levels of protein abundances. For instance, AlsS was much more abundant than CATec3 Y20F in the *alsS-cat* operon, IlvC more abundant than YqhD in the *ilvC-ilvD-yqhD* operon, and LeuA* more abundant than LeuB, LeuC, and LeuD in the *leuABCD* operon (Figs. 2d, 3c).

Globally, we observed a large perturbation in the amino acid reallocation constituting the proteome. The mass fraction contributions of cysteine (+27%), methionine (+10%), serine (+7%), glycine (+5%), aspartate (+3.2%) and alanine (+2.3%) in the proteome increased at least 2% while those of tryptophan (−13%), valine (−5.2%), threonine (−3.8%), glutamate (−3.8%), proline (−3.5%), tyrosine (−2.5%), phenylalanine (−2.5%), arginine (−2.2%), and histidine (−2.1%) decreased at least 2% (Fig. 3e). This analysis was based on the aggregate abundance for isoamyl acetate pathway proteins and their amino acid distributions relative to those observed in the quantifiable proteome. Since the isoamyl acetate pathway proteins represent an outsized share of protein abundance upon induction, the resource demand of the pathway module could impart a sizable burden on the rest of the system (i.e., amino acid biosynthesis, transcription and translation machinery, energetics, etc.)

Taken together, overexpression of the isoamyl acetate pathway and its effect on proteome allocation suggest an imposed metabolic burden. The severe fold-change abundance difference of CATec3 Y20F upon induction, as compared to AlsS, and high accumulation of isoamyl alcohol implied that CATec3 Y20F might be the rate limiting step.

#### Tuning the isoamyl acetate pathway by enhancing CATec3 Y20F expression

To test whether the AAT activity was the bottleneck for the isoamyl acetate production, we introduced an additional monocistronic CATec3 Y20F gene under the control of T7lac promoter on the medium copy pCDF plasmid (Fig. 4a). The additional expression of CATec3 Y20F in HSEC1311 did not affect cell growth as compared to HSEC1210 (Fig. 4b) but improved isoamyl acetate production by 4.3-fold, reaching a titer of 3.1 g/L (Fig. 4c). Isoamyl alcohol accumulation in HSEC1311 was significantly reduced by 10-fold from 2.93 g/L to 0.22 g/L (Fig. 4d). This perturbation reduced the enzyme aggregate abundance of the isoamyl acetate pathway from 31.9% to 22.6%, while the reallocation for generation of precursor metabolites and energy and amino acids were enhanced (Figs. 4g, S2). The CATec3 Y20F abundance was 10.2-folds higher in HSEC1311 than HSEC1210 while other proteins of the isoamyl acetate pathway exhibited a 1.7- to 3-folds decrease in protein abundance (Fig. 4f), likely due to the transcriptional and/or translational competition by the introduction of an additional CATec3 Y20F operon. Remarkably, the additional expression of CATec3 Y20F resulted in the amino acid reallocation in the global proteome (Fig. 4e). Mass fractions of some amino acids in the proteome became more abundant in HSEC1311 than HSEC1210 while others were reduced. The trend observed here is reciprocal to the scenario of HSEC1210 growing in the media with and without IPTG induction. The mass fraction contributions of tryptophan (+7.0%), tyrosine (+3.1%), and phenylalanine (+2.7%) in the proteome increased at least 2% while those of cystine (−6.7%), serine (−2.9%), and methionine (−2.5%) decreased at least 2 %.

**Figure 4.**
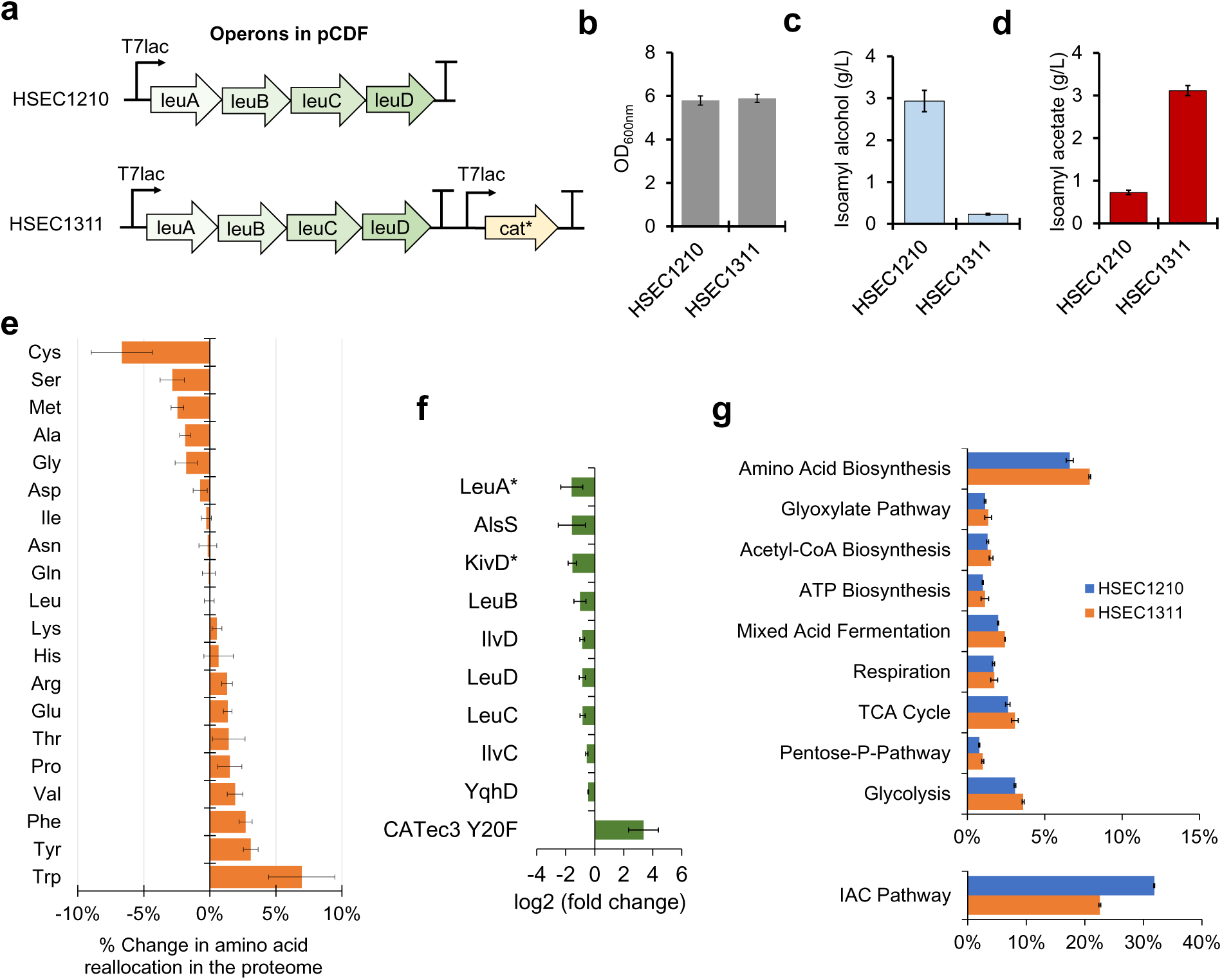
Alleviation of metabolic bottleneck and burden of the isoamyl acetate pathway by overexpression of CATec3 Y20F. **(a)** Genetic architecture of operons in the pCDFDuet-1 backbone plasmid of HSEC1210 and HSEC1311. In addition to having CATec3 Y20F in the pRSF plasmid like HSEC1210, HSEC1311 was designed to express additional CATec3 Y20F in the pCDF plasmid. **(b-d)** Comparison of cell growth **(b)** and production of isoamyl alcohol **(c)** and isoamyl acetate **(d)** by HSEC1210 and HSEC1311 after 48 h. **(e)** Percent of change in amino acid reallocation in the proteome upon additional expression of CATec3 Y20F. **(f)** Comparison of proteomic fold-changes of enzymes in the isoamyl acetate pathway between HSEC1210 and HSEC1311 at 24 h. **(g)** Mass fractions of the pathways for generating precursor metabolites and energy and biosynthesis pathways of amino acids in the proteomes. Each data represents mean ± 1 standard deviation from at least three biological replicates.

Overall, CATec3 Y20F was the rate limiting step in HSEC1210. The additional overexpression of this protein in HSEC1311 helped alleviate both the metabolic bottleneck and metabolic burden.

### Demonstration of high-level isoamyl acetate production

#### Further deletion of upstream competing pathways did not improve isoamyl acetate production

We next examined whether additional deletion of competitive upstream pathways in HSEC1311 could further improve production of isoamyl acetate. Our deletion targets included *ldhA* (L-lactate dehydrogenase), *ackA-pta* (an operon of acetate kinase and phosphate acetyltransferase), *ilvE* (branched chain amino acid aminotransferase), and *tyrB* (aromatic amino-acid aminotransferase), which can potentially help reduce the lactate and acetate formation as byproducts and improve ketoacid availability (Fig. 5a). The isoamyl acetate production modules were plugged into the engineered strains and isoamyl acetate yields were compared (Figs. 5b, S3). Our results showed that deletion of the upstream pathway enzymes did not significantly improve isoamyl acetate yield, suggesting that the upstream competitive pathways were not major metabolic bottlenecks in our system. Isoamyl acetate yield reached up to 0.17 (g/g glucose) that corresponds to 47% of the maximum theoretical yield (0.36 g/g).

**Figure 5.**
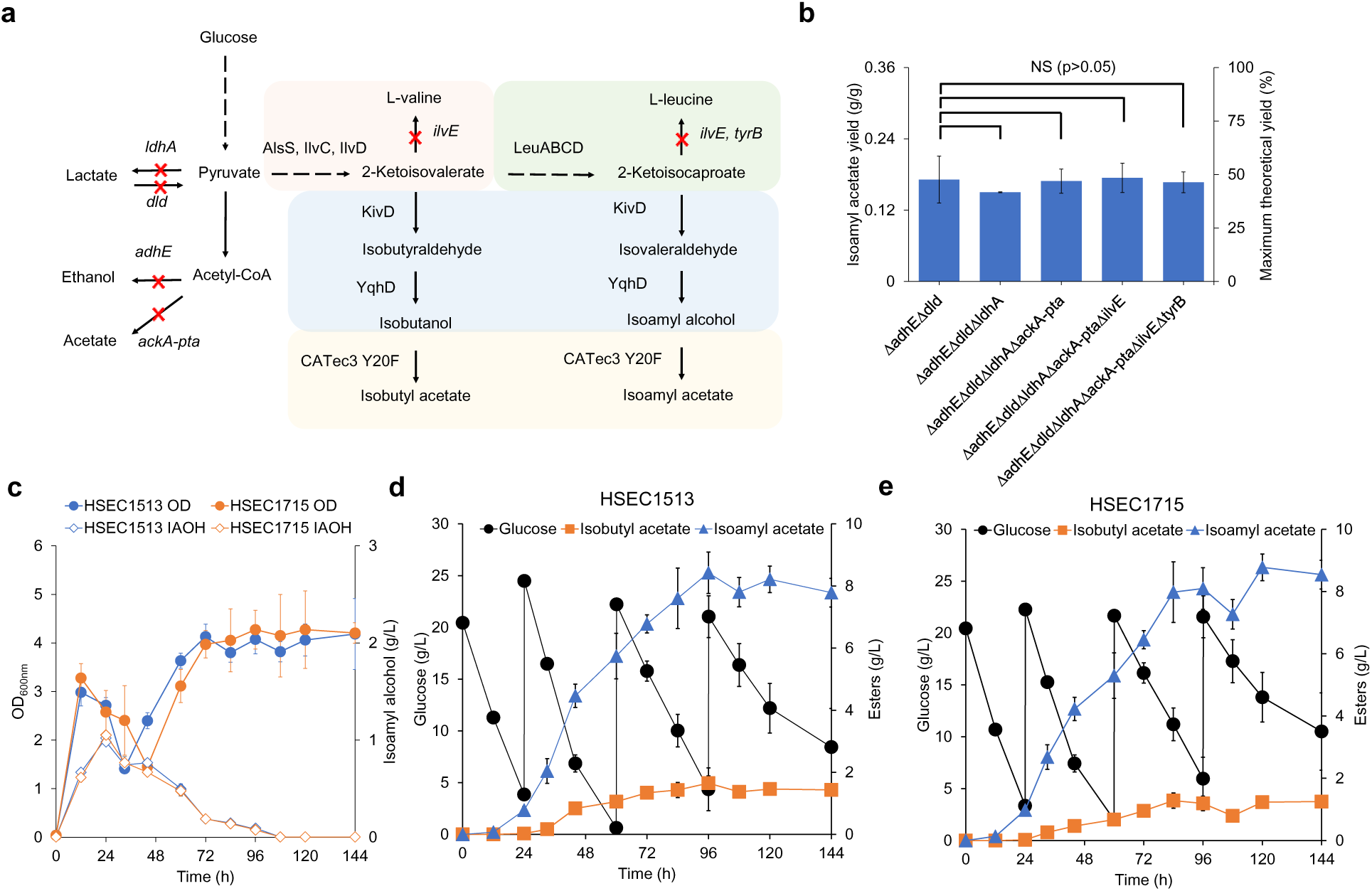
Optimization of isoamyl acetate production. **(a)** Metabolic map displaying deletion of genes participating in pathways that might compete for isoamyl acetate biosynthesis. The red marks indicate gene deletions and their reactions. **(b)** Comparison of isoamyl acetate yield on glucose between the engineered strains. **(c)** Profiles of cell growth and accumulated isoamyl alcohol production during fed-batch culture by HSEC1412 and HSEC1614. **(d-e)** Kinetics of glucose consumption, isobutyl acetate production, and isoamyl acetate production from fed-batch culture by **(d)** HSEC1412 and **(e)** HSEC1614. Each data represents mean ± 1 standard deviation from at least three biological replicates.

#### Fed-batch fermentation boosted high-level production of isoamyl acetate

The considerable isoamyl acetate yield prompted us to investigate whether glucose-fed batch fermentation with pH control could increase isoamyl acetate production. Because the *ilvE* and *tyrB* deletions can affect amino acid utilization (Iwasaki et al., 2021) during a prolonged isoamyl acetate production, we characterized and compared the isoamyl acetate production of the two engineered strains, including HSEC1513 (BL21 (DE3) Δ*adhE* Δ*dld* Δ*ldhA* Δ*ackA-pta* harboring the isoamyl acetate production modules) and HSEC1715 (BL21 (DE3) Δ*adhE* Δ*dld* Δ*ldhA* Δ*ackA-pta* Δ*ilvE* Δ*tyrB* harboring the isoamyl acetate production modules) over 144 h (Figs. 5c, 5d, 5e). Remarkably, we observed accumulation of isoamyl alcohol from both strains for the first 24 h and optical density (OD_600nm_) decreased from 3.2 to 1.4 between 12 to 36 h, probably due to alcohol toxicity (Fig. 5c). The OD_600nm_ later increased up to 4.3, and the accumulated isoamyl alcohol was converted to isoamyl acetate for the next 72 h. HSEC1513 and HSEC1715 produced isoamyl acetate up to 8.4 g/L (82% selectivity) and 8.8 g/L (86% selectivity), respectively, reporting the highest microbial production of isoamyl acetate titers up to date (Figs. 5d, 5e). The byproduct isobutyl acetate was produced up to 1.4 g/L. Since HSEC1513 and HSEC1715 showed little difference in cell growth and isoamyl acetate production, we concluded that the aminotransferases were not the major bottlenecks in our engineered strains.

## DISCUSSION

The one-carbon recursive elongation pathway is important for making branched-chain amino acids, aldehydes, alcohols, and esters. Due to this complex and highly branched pathway, controlling selective microbial biosynthesis of these target molecules has been an outstanding metabolic engineering problem. To address the problem, we developed a generalizable modular design framework to systematically tune selective microbial biosynthesis of branched-chain acetate esters that require the integration of four submodules including the KIV submodule, the elongation submodule, the KDC submodule, and the AAT submodule. We validated this framework by demonstrating selective biosynthesis of isoamyl acetate over isobutyl acetate as an ester byproduct, achieving the highest titer (8.7 g/L), yield (0.17 g/g), and selectivity (86%) reported to date.

Critical to selective microbial biosynthesis of branched-chain acetate esters is control of protein expression and specificity of the first committed steps of the four submodules including AlsS, LeuA*, Kivd*, and CAT*. To achieve high production of isoamyl acetate, the engineered pathway required overexpression of 10 genes, which made up about 23-31% of total proteome allocation and represented a relatively large fractional share of overall proteome abundance as compared to the non-overexpressed control. This metabolic pathway rewiring caused global perturbations that can be seen in the fraction of amino acids within the proteome. This metabolic tradeoff occurred at the expense of other cellular processes such as the fueling pathways responsible for generating precursor metabolites, and energy and amino acid biosynthesis, which could explain the observed metabolic burden affecting cell growth and yield. While the push-and-pull strategy of metabolic fluxes towards the target pathway(s) is commonly practiced in metabolic engineering, our study provides direct quantitative evidence of the proteome reallocation required to achieve pathway efficiency and potential metabolic tradeoffs.

Remarkably, close examination of the engineered pathway proteins shows that increase in abundances of target proteins might be significantly different even though their encoding genes are organized in the same operon. Upstream genes exhibited larger relative increases in protein abundance relative to downstream ones within an operon. Different amino acid requirements, codon usage, and/or protein folding efficiency for each protein might have contributed to this discrepancy. Our result further revealed that a single limiting enzymatic step, such as AAT, could impose a detrimental metabolic bottleneck/burden due to flux imbalance and hence accumulation of alcohol intermediates that become inhibitory.

Medium chain length (C_6_-C_10_) branched esters are relatively hydrophobic metabolites and therefore, toxic to cells as they interfere with cell membranes (Wilbanks and Trinh, 2017a). Due to low solubility of these esters in aqueous solutions, our study demonstrated the feasibility of producing them at much higher concentrations than their reported toxicity limit (0.25 g/L) via *in situ* fermentation and extraction. Our data also demonstrate that ester biosynthesis could help detoxify alcohols as intermediates via overexpression of AAT whereby the resulting esters are immediately extracted by a non-toxic solvent overlay such as hexadecane. Even though the enhanced expression level of CATs improved isoamyl acetate production, accumulation of isoamyl alcohol and decrease in cell growth observed during the fed-batch fermentation suggested that the AAT activity was still a major bottleneck (Fig. 5c).

Unlike other eukaryotic AATs, use of CATec3 Y20F is beneficial for designer ester production due to higher solubility, thermostability, and selectivity (Seo et al., 2021). However, it has relatively lower catalytic efficiency towards short chain alcohols such as isobutanol and isoamyl alcohol. Thus, future protein engineering of CATec3 Y20F for improved catalytic efficiency towards isoamyl alcohol can help reduce the alcohol accumulation while not requiring high protein expression. Although our study focused on the selective microbial biosynthesis of isoamyl acetate as a proof-of-concept, the generalizable modular design of the recursive one-carbon elongation pathway can be extended to produce longer branched-chain acetate esters such as isohexyl acetate and isoheptyl acetate. Since CATec3 Y20F has higher catalytic efficiency towards longer chain alcohols (Seo et al., 2021), it is expected that the selectivity of KDC and/or LeuA needs to be further engineered to achieve selective microbial biosynthesis of designer acetate esters with longer carbon chain lengths.

One potential challenge of branched chain acetate ester biosynthesis is the stoichiometric redox imbalance that might cause growth inhibition and/or impaired production. Fermentative isoamyl acetate production requires two moles of glucose to produce two acetyl-CoAs and one isoamyl alcohol (2 Glucose + 5 NAD(P)+ = 1 Isoamyl acetate + 5CO_2_ + 5NAD(P)H). Therefore, the pathway generates five moles of excess NAD(P)H that could inhibit the fermentation under an oxygen limited condition without an appropriately coupled electron sink(s). Indeed, HSEC1311 was not able to grow without oxygen (Fig. S4), suggesting the possible inhibition of fermentation by the cofactor imbalance under the oxygen limited conditions. Recent studies suggest that modular cell design principle can harness such electron redundancy for multi-objective strain design by coupling production pathways with cell growth (Garcia and Trinh, 2019; Wilbanks et al., 2018). Therefore, further optimization should include process engineering and chassis cell design for improved production of branched-chain acetate esters.

## MATERIALS AND METHODS

### Strains and plasmids

*E. coli* DH5α and BL21(DE3) were used for molecular cloning and ester production, respectively. The strains and plasmids used are listed in Table 1.

**Table 1:**
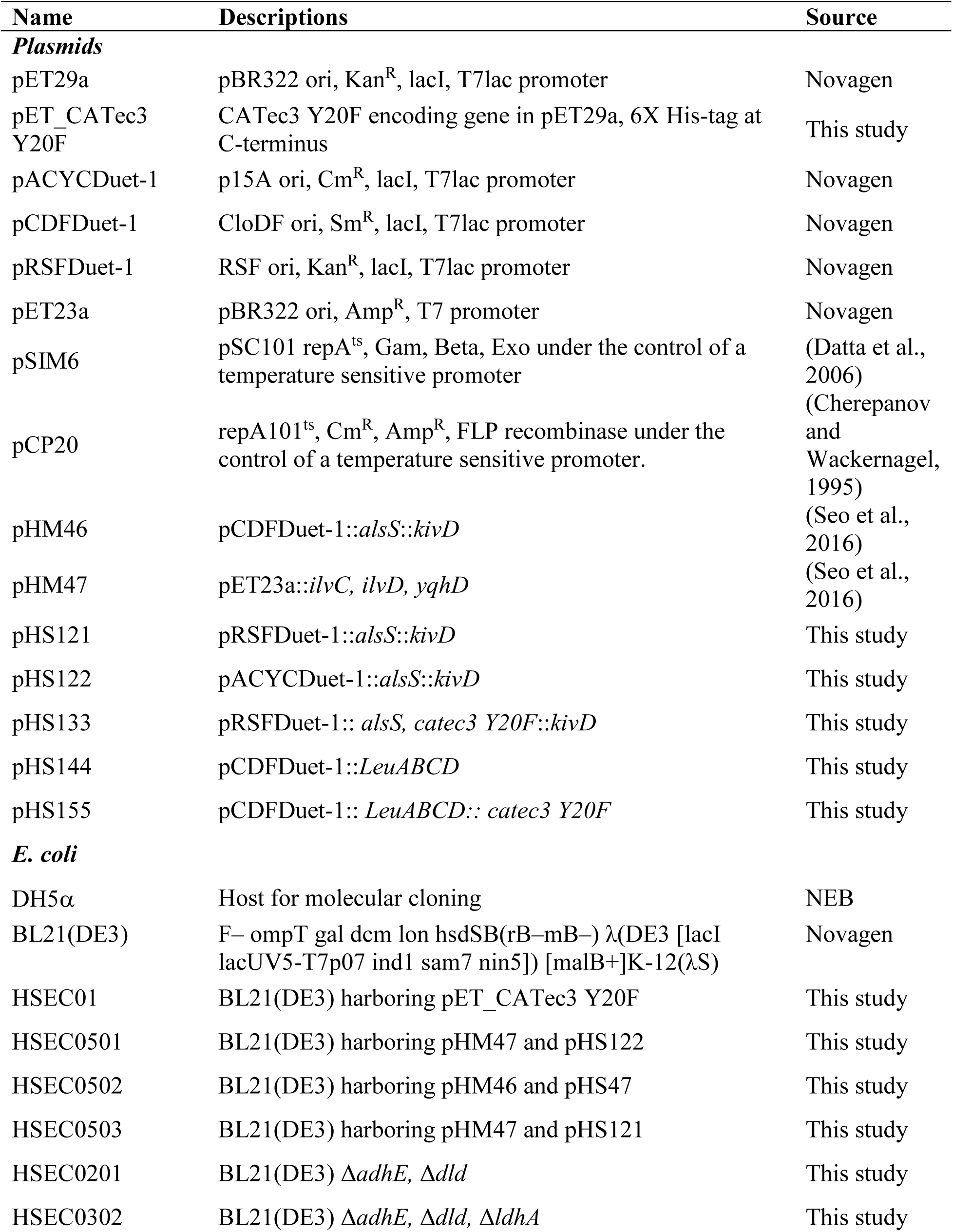
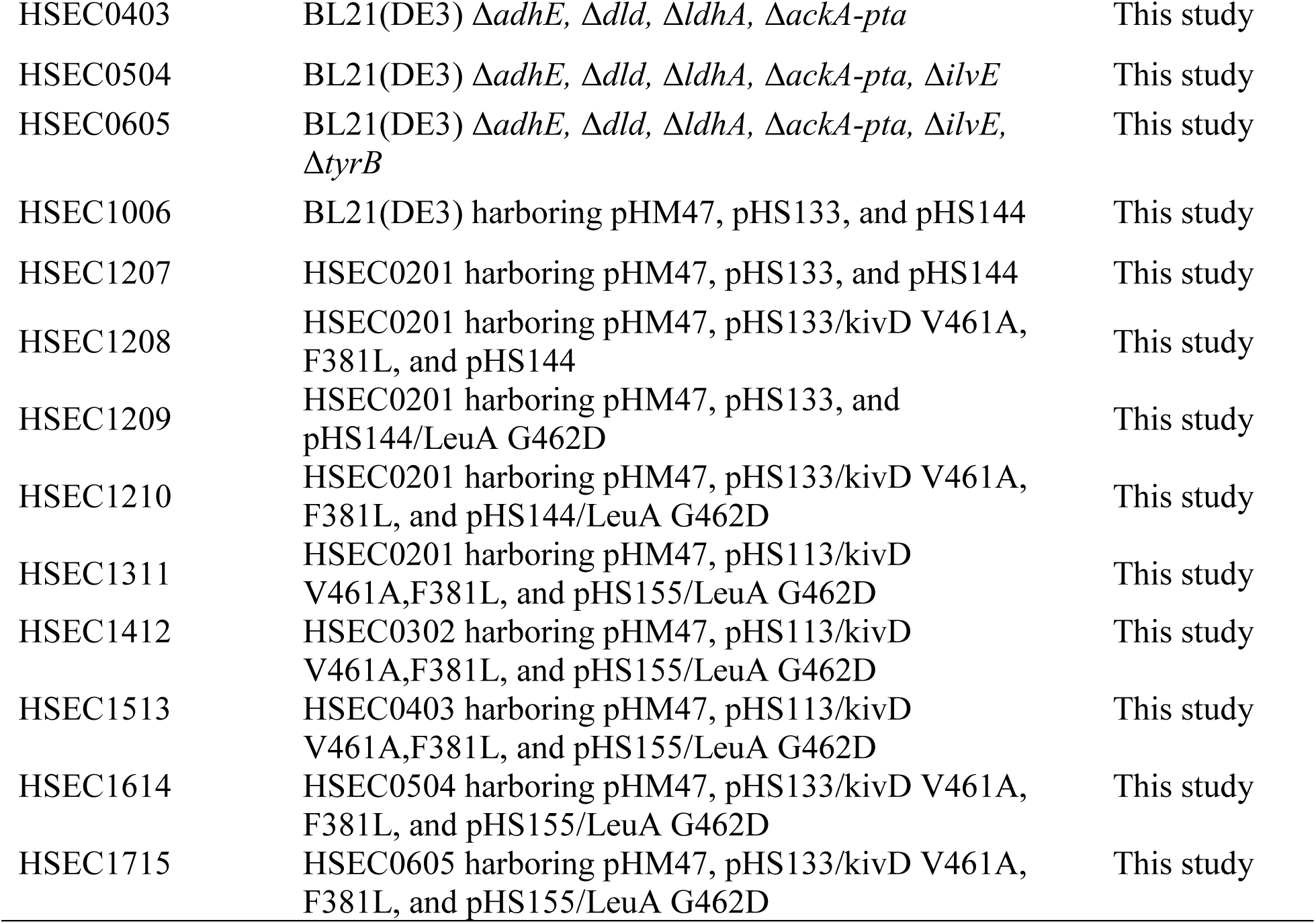
A list of plasmids and strains used in this study.

### Media and cultivation

*E. coli* strains were grown in lysogeny broth (LB) medium or M9 hybrid medium containing glucose as a carbon source and 5 g/L yeast extract supplemented with 100 µg/mL ampicillin and/or 100 μg/mL spectinomycin and/or 50 µg/mL kanamycin when appropriate.

For the batch fermentation, cells were cultured microaerobically in a 125 mL screw-capped shake flask with a working volume of 20 mL in M9 medium containing 30 g/L glucose and 5 g/L yeast extract. 10 mL of hexadecane (50% v/v) was overlaid to extract esters produced during the fermentation. To compare performance of the engineered strains, cells were cultured at 37℃ for 48 h and the culture supernatant and hexadecane layer were analyzed. For the fed-batch fermentation, 20g/L glucose working concentration was intermittently added to the culture using 600 g/L glucose stock when the glucose concentration was below 5 g/L. The pH was adjusted with 5M KOH to maintain its value between 6.0 and 7.5 every 12 h after 24 h until 120 h.

### Molecular cloning

#### Plasmid construction

Plasmids were constructed by ligation-dependent cloning and/or Gibson DNA assembly. Briefly, DNA fragments were amplified using the Phusion DNA polymerase (cat# F530S, Thermo Fisher Scientific, MA, USA) and then purified by DNA purification and gel extraction kits (Omega BioTek, GA, USA). For the ligation-dependent cloning, the vectors and inserts were digested by restriction enzymes and ligated together using a T4 DNA ligase. In the case of Gibson DNA assembly cloning, the purified DNA fragments of the vector and insert were mixed together with the Gibson master mix (Gibson et al., 2009) and assembled at 50℃ for 1 h. Using the DNA mixtures, *E. coli* DH5α was transformed by heat-shock transformation and selected on LB agar plates (15 g/L agar) with appropriate antibiotics. All the constructed plasmids were checked by PCR amplification and/or restriction enzyme digestion, and/or Sanger sequencing. The primers used in this study are listed in Table S1.

#### Recombineering

*E. coli* gene deletions were carried out using recombineering (Sharan et al., 2009). A temperature sensitive low-copy plasmid that contains exo, bet, and gam in their native phage operon, pL, under λ CI repressor control (pSIM6) was used to induce homologous recombination of double strand DNA into the genome (Datta et al., 2006). Briefly, *E. coli* strains harboring pSIM6 was cultured in 3 mL LB medium at 30℃ overnight. The grown cells were transferred to fresh 20 mL LB medium in a 250 mL flask with 1% inoculum size (200 μL) and cultured for 2-3 hours in a water bath shaking incubator at 200 rpm and 32℃. At OD_600nm_ of 0.4∼0.6, the cell culture flask was transferred to a preheated 42℃ water bath shaking incubator at 200 rpm for 15 minutes (min). Then, the cells were immediately cooled down in ice for 10 mins and centrifuged at 4,700 rpm for 10 min. The cell pellets were washed twice with 50 mL ice-cool sterile Millipore water and then suspended in the 200 μL ice-cool sterile Millipore water. 80 mL of the concentrated cells were mixed with ∼100 ng of linear double-stranded DNA containing FRT-Kan-FRT cassette, amplified by PCR. Then, the cells were transferred to an ice-chilled 1- mm gap electroporation cuvette (BTX Harvard Apparatus, MA, USA) followed by an exponential decay pulse with 1.8 kV, 350 Ω, and 25 μF, which gave usual pulse duration of 4.5-6.0 ms. The cells were immediately mixed with 700 mL LB medium and recovered in a shaking incubator at 30℃ for 2 h. The recovered cells were plated on LB solid medium with 25 mg/mL kanamycin and incubated at 30℃ for 2 days. Successful gene deletion was confirmed by colony PCR using multiple combinations of primers specifically binding at upstream and/or downstream of the target location and/or Kan in the FRT-Kan-FRT cassette (Table S1). The kanamycin resistance marker was subsequently disrupted by FLP mediated recombination of FRT by pCP20 (Datsenko and Wanner, 2000).

### Analytical methods

#### Cell growth measurement

Cell growth was measured by optical density (OD) with a spectrophotometer (Spectronic 200+, Thermo Fisher Scientific, MA, USA) and/or a microplate reader (Synergy HTX microplate reader, BioTek) at 600 nm wavelength.

#### 3,5-dinitrosalicylic *acid (DNS) assay*

Slightly modified DNS method was used to quickly quantify and monitor the glucose consumption during the glucose fed-batch culture (Miller, 1959). Briefly, 10 μL of 1-, 2-, and 4-times diluted culture supernatants were mixed with 200 μL of the DNS reagent consisting of 16 g/L NaOH, 5 g/L phenol, 5 g/L sodium sulfite, and 300 g/L potassium sodium tartrate, and 10 g/L DNS, and incubated at 98°C for 10 mins. The samples were read by a microplate reader at 540nm. The M9 medium with 0 g/L, 1 g/L, 3 g/L, 5 g/L, and 10 g/L glucose concentration was used as standard for every reaction.

#### Proteomics

Engineered *E. coli* strains (HSEC1210 uninduced; HSEC1210 induced with IPTG; HSEC1311 induced with IPTG) were cultured as described above and harvested at 24 h in biological triplicates. Cells were pelleted, supernatants removed, and pellets snap frozen and stored at -80°C. Frozen pellets (∼100 µL pellet volume) were then processed for LC-MS/MS-based proteomic measurements by resuspending in cold 100 mM Tris-HCl buffer, pH 8.0, adding ∼200 µL of 0.15 zirconium oxide beads, and bead beating with a Geno/Grinder 2010 (SPEX SamplePrep) for 5 min at high speed (1,750 rpm). Crude cell protein lysates were then further processed and proteolytically digested with trypsin as previously described (Walker et al., 2021). Peptide samples were analyzed by automated 1D LC-MS/MS analysis using a Vanquish UHPLC plumbed directly to a Q Exactive Plus mass spectrometer (Thermo Scientific) outfitted with a trapping column coupled to an in-house pulled nanospray emitter (Walker et al., 2021). For each sample, 3 µg of peptides were loaded, desalted, and separated by uHPLC with the following conditions: sample injection followed by 100% solvent A (95% H_2_O, 5% acetonitrile, 0.1% formic acid) chase from 0-30 min (load and desalt), linear gradient 0% to 30% solvent B (70% acetonitrile, 30% water, 0.1% formic acid) from 30-220 min (separation), and column re-equilibration at 100% solvent A from 220-240 min. Eluting peptides were measured and sequenced by data-dependent acquisition on the Q Exactive MS as previously described (Clarkson et al., 2017).

#### High-performance liquid chromatography (HPLC) analysis

Extracellular metabolites were quantified using HPLC system (Shimadzu Inc., MD, USA). 800 μL of culture samples were centrifuged at 17,000 xg for 3 m followed by filtering through a 96-well filter plate with 0.45 micron. The samples were run with 10 mM sulfuric acid at 0.6 mL/min flow rate on an Aminex HPX-87H (Biorad Inc., CA, USA) column at 50°C. Concentrations of sugars, organic acids, and alcohols were determined by refractive index detector (RID) and ultra-violet detector (UVD).

#### Gas chromatography coupled with mass spectroscopy (GC/MS) analysis

Esters were quantified by GC (HP 6890, Agilent, CA, USA) equipped with a MS (HP 5973, Agilent, CA, USA). A Zebron ZB-5 (Phenomenex, CA, USA) capillary column (30 m x 0.25 mm x 0.25 μm) was used with helium as the carrier gas at a flow rate of 0.5 mL/min. The oven temperature was programed as follows: 50°C initial temperature, 1°C/min ramp up to 58°C, 25°C/min ramp up to 235°C, 50°C/min ramp up to 300°C, and 2-minutes bake-out at 300°C. 1 μL sample was injected into the column with the 1:50 split mode at an injector temperature of 280°C. For the MS system, selected ion mode (SIM) was used to detect and quantify esters with the parameters described previously(Seo et al., 2021). As an internal standard, 10 mg/L n-decane were added in initial hexadecane layer and detected with m/z 85, 99, and 113 from 12 to 15 m retention time range.

### Computational analysis

#### Proteomic analysis

MS/MS spectra were searched against the *E. coli* BL21(DE3) proteome (UniProt; Aug21 build) appended with relevant exogenous protein sequences and common protein contaminants using the MS Amanda v.2.0 algorithm in Proteome Discoverer v.2.3 (Thermo Scientific). As described previously(Walker et al., 2021), peptide spectrum matches (PSM) were required to be fully tryptic with up to 2 miscleavages; a static modification of 57.0214 Da on cysteine (carbamidomethylated) and a dynamic modification of 15.9949 Da on methionine (oxidized) residues. PSMs were scored and filtered using Percolator and false-discovery rates initially controlled at < 1% at both the PSM- and peptide-levels. Peptides were then quantified by chromatographic area-under-the-curve, mapped to their respective proteins, and areas summed to estimate protein-level abundance. Protein abundances were log2 transformed, and sample abundance distributions normalized by LOESS then median centered in InfernoRDN (Taverner et al., 2012). Missing values were imputed and sample groups statistically assessed in Perseus(Tyanova et al., 2016).

All raw mass spectra for quantification of proteins used in this study have been deposited in the MassIVE and ProteomeXchange data repositories under accession numbers MSV000088838 (MassIVE) and PXD031694 (ProteomeXchange), with data files available at ftp://massive.ucsd.edu/MSV000088838/.

#### Proteome reallocation analysis

EcoCyc version 25.5 was used to extract annotated genes, proteins, and pathways for proteome mapping of core metabolism using *E. coli* BL21(DE3) (Caspi et al.). These pathways include i) generation of metabolites and precursors (glycolysis, pentose phosphate pathway, Krebs cycle, glyoxylate cycle, respiration, fermentation, ATP biosynthesis, acetyl CoA biosynthesis, and Entner Doudoroff pathway), ii) amino acid biosynthesis, iii) aminoacyl-tRNA charging, and iv) isoamyl acetate biosynthesis. Amino acid sequences of each protein in the proteome were retrieved from UniProt (UniProt, 2021).

Mass fraction of protein P_i_, f_Pi_, in the proteome is calculated as follows:

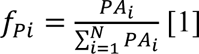

where PA_i_ is the abundance of protein i in the proteome, N is the total number of proteins in the proteome measured, and 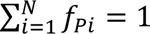. For a metabolic pathway with M proteins, the mass fraction of pathway proteome, f_Path_, is determined as follows:

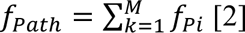

Mass fraction of amino acid j, f_Aj_, in the proteome is calculated as follows:

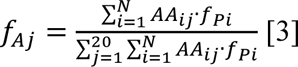

where AA_ij_ is the abundance of amino acid j in protein i of the proteome and 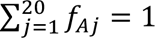.

## ACKNOWLEDGMENTS

This research was financially supported in part by the DOE BER award (DE-SC0022226) and the DOE subcontract grant (DE-AC05-000R22725) by the Center of Bioenergy Innovation, the U.S. Department of Energy Bioenergy Research Center funded by the Office of Biological and Environmental Research in the DOE Office of Science, and the U.S. Department of Energy Joint Genome Institute. The work conducted by the U.S. Department of Energy Joint Genome Institute, a DOE Office of Science User Facility, is supported under Contract No. DE-AC02-05CH11231. The authors would like to thank the Center of Environmental Biotechnology at UTK for using the GC/MS instrument.

## CREDIT AUTHOR STATEMENT

**Hyeongmin Seo:** Conceptualization, Methodology, Validation, Formal analysis, Investigation, Data Curation, Writing-Original Draft, Visualization. **Richard J. Giannone:** Methodology, Validation, Formal analysis, Investigation, Data Curation, Writing-Review & Editing. **Yung-Hun Yang:** Resources, Writing-Review & Editing. **Cong T. Trinh:** Conceptualization, Methodology, Formal analysis, Investigation, Data Curation, Supervision, Project administration, Funding acquisition, Writing-Original Draft, Review & Editing.

## SUPPLEMENTARY DATA

**Supplementary File S1** contains Table S1 and Figures S1, S2, S3, S4.

**Table S1:** A list of primers used in this study.

**Figure S1**. Relative catalytic efficiency of AATs towards isoamyl alcohol over isobutanol.

**Figure S2.** Comparative proteomics of HSEC1311 and HSEC1210 growing in media with IPTG induction at 24h.

**Figure S3**. Isoamyl acetate titers of engineered strains with sequential gene deletions after 48 h culturing.

**Figure S4.** Growth of HSEC1311 under anaerobic and microaerobic conditions.

