## Supplementary File S1 for "Proteome reallocation enables the selective *de novo* biosynthesis of non-linear, branched-chain acetate esters"

### Proteome reallocation enables selective *de novo* biosynthesis of non-linear, branched-chain acetate esters

Hyeongmin Seo[1,2], Richard J. Giannone[2,3], Yung-Hun Yang[4], and Cong T. Trinh[1,2§]

[1]Department of Chemical and Biomolecular Engineering, The University of Tennessee, Knoxville, TN, USA

[2]Center of Bioenergy Innovation, Oak Ridge National Laboratory, Oak Ridge, TN, USA

[3]Chemical Sciences Division, Oak Ridge National Laboratory, Oak Ridge, TN 37830, USA

[4]Department of Biological Engineering, Konkuk University, Seoul, Republic of Korea

**Table S1.** A list of primers used in this study. The bold and underlined letters indicate restriction and site-directed mutation sites, respectively.

| Primer name | Primer sequence (5' to 3') | Description |
|---|---|---|
| HS677 | CTCT**GTCGAC**TTAAGAAGGAGATA TAGATATGAATTATACTAA | CATec3 Y20F cloning forward (pHS133 construct) |
| HS678 | CTCTAA**GCGGCCGC**TTACTTCAATT TCGAATTGCAGAGC | CATec3 Y20F cloning reverse (pHS133 construct) |
| HS683 | CTCT**GAGCTC**TATGAGCCAGCAAG TCATTATTTTCGATAC | LeuABCD cloning forward (pHS144 construct) |
| HS684 | CTCTAA**CTCGAG**TTAATTCATAAAC GCAGGTTGTTTTGC | LeuABCD cloning reverse (pHS144 construct) |
| HS802 | CTCTAA**GCGGCCGC**TTAATTCATA AACGCAGGTTGTTTTGC | LeuABCD cloning at MCS1 reverse (pHS155 construct) |
| HS763 | CATATGTATATCTCCTTCTTATACTT AACTAA | Backbone amplification forward for CATec3 Y20F cloning (pHS155 construct) |
| HS764 | ATTAACCTAGGCTGCTGCCACCGCT GAGCAA | Backbone amplification reverse for CATec3 Y20F cloning (pHS155 construct) |
| HS785 | AGTATAAGAAGGAGATATACATAT GAATTATACTAAATTCGATG | CATec3 Y20F cloning forward (pHS155 construct) |
| HS786 | TTGCTCAGCGGTGGCAGCAGCCTAG GTTAATTTACTTCAATTTCGAATTGC | CATec3 Y20F cloning reverse (pHS155 construct) |
| HS701 | ATGCGCTG GAC CAGGTGGATATCGTCGC | LeuA G462D mutation forward |
| HS702 | CACCTG GTC CAGCGCATCTTTACCGTGG | LeuA G462D mutation reverse |
| HS703 | TTATACA GCG GAAAGAGAAATTCATGG | kivD V461A mutation forward |
| HS704 | TCTTTC CGC TGTATAACCATCATTATTG | kivD V461A mutation reverse |
| HS705 | GGACATCA CTG TTTGGCGCTTCATCA | kivD F381L mutation forward |
| HS706 | CGCCAAA CAG TGATGTCCCTTGTTCAG | kivD F381L mutation reverse |
| HS33 | CGTTGGCTACCCGTGATATT | FRT-Kan-FRT cassette forward for deletion confirmation |
| HS34 | GCCCAGTCATAGCCGAATAG | FRT-Kan-FRT cassette reverse for deletion confirmation |
| HS117 | ATTTACTAAAAAAGTTTAACATTAT CAGGAGAGCATTATG GTGTAGGCTGGAGCTGCTTC | adhE deletion forward |
| HS118 | GCCCAGAAGGGGCCGTTTATGTTGC CAGACAGCGCTACTGA CATATGAATATCCTCCTTA | adhE deletion reverse |
| HS372 | CTATACTCTCGTATTCGAGCAGATG | adhE deletion check upstream |
| HS373 | GGCCGTTTATATTGCCAGACAG | adhE deletion check downstream |

| | | |
|---|---|---|
| HS31 | TTTGTGATATTTTTTCGCCACCACAAGGAGTGGAAAATGGTGTAGGCTGGAGCTGCTTC | dld deletion forward |
| HS32 | TAAGTGAATTCGGATGGCGATACTCTGCCATCCGTAATTTCATATGAATATCCTCCTTA | dld deletion reverse |
| HS39 | CAGTTTATTGTCTGAATTTTCAAAATA | dld deletion check upstream |
| HS40 | AGCTATAAAAAACAAAAAGCCGC | dld deletion check downstream |
| HS863 | GCTTAAATGTGATTCAACATCACTGGAGAAAGTCTTATGGTGTAGGCTGGAGCTGCTTC | ldhA deletion forward |
| HS864 | TCCCCTGCAACCCAGGGGAGCTGATTCAGATAATCCCCAATCATATGAATATCCTCCTTA | ldhA deletion reverse |
| HS865 | CCCGAGCGTCATCAGCAGCG | ldhA deletion check upstream |
| HS866 | GGTCATTGCCAGCCCTTTGGCTG | ldhA deletion check downstream |
| HS922 | ATGGCTCCCTGACGTTTTTTTAGCCACGTATCAATTATAGGTACTTCCATGGTGTAGGCTGGAGCTGCTTC | ackA-pta deletion forward |
| HS923 | ATTATTTCCGGTTCAGATATCCGCAGCGCAAAGCTGCGGATGATGACGAGACATATGAATATCCTCCTTA | ackA-pta deletion reverse |
| HS924 | ACGCAAAATGGCATAGACTCAAGATAT | ackA-pta deletion check upstream |
| HS925 | CACAAAACAAAGTGGTAAGTATGCAAAGT | ackA-pta deletion check downstream |
| HS713 | CAAATCCGCGCCTGAGCGCAAAAGGAATATAAAAATGGTGTAGGCTGGAGCTGCTTC | ilvE deletion forward |
| HS714 | AATGGGACGGTGCGTGCCGTCCCATTTTTTGTATCATATGAATATCCTCCTTA | ilvE deletion reverse |
| HS717 | AGTCAGTTAAATAAACTG | ilvE deletion check upstream |
| HS718 | GCCATGGGTGGTGGTGGC | ilvE deletion check downstream |
| HS715 | TAACCACCTGCCCGTAAACCTGGAGAACCATCGCGTGGTGTAGGCTGGAGCTGCTTC | tyrB deletion forward |
| HS716 | GCTGGGTAGCTCCAGCCTGCTTTCCTGCATTACACATATGAATATCCTCCTTA | tyrB deletion reverse |
| HS720 | GTTGCTAATTGCCGTTC | tyrB deletion check upstream |
| HS873 | TAGAACGATGGCATCAAAA | tyrB deletion check downstream |

**Figure S1.** Relative catalytic efficiency of AATs towards isoamyl alcohol over isobutanol. The catalytic efficiencies of ATF1, ATF2, LuxE, BPBT, and SAAT were reported by Tai *et al.* (Tai et al., 2015) while the catalytic efficiency of CATec3 Y20F was investigated by Seo *et al.* (Seo et al., 2021).

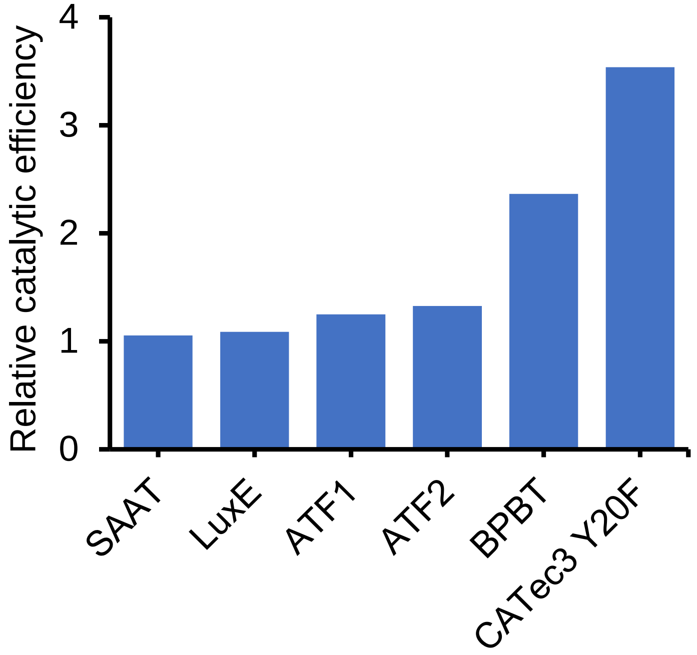

**Figure S2.** Comparative proteomics of HSEC1311 and HSEC1210 growing in media with IPTG induction at 24h. The data represent mean values from three biological replicates. The pval and diff values are referred to -$\log_{10}$(p-value) and $\log_2$(difference), respectively.

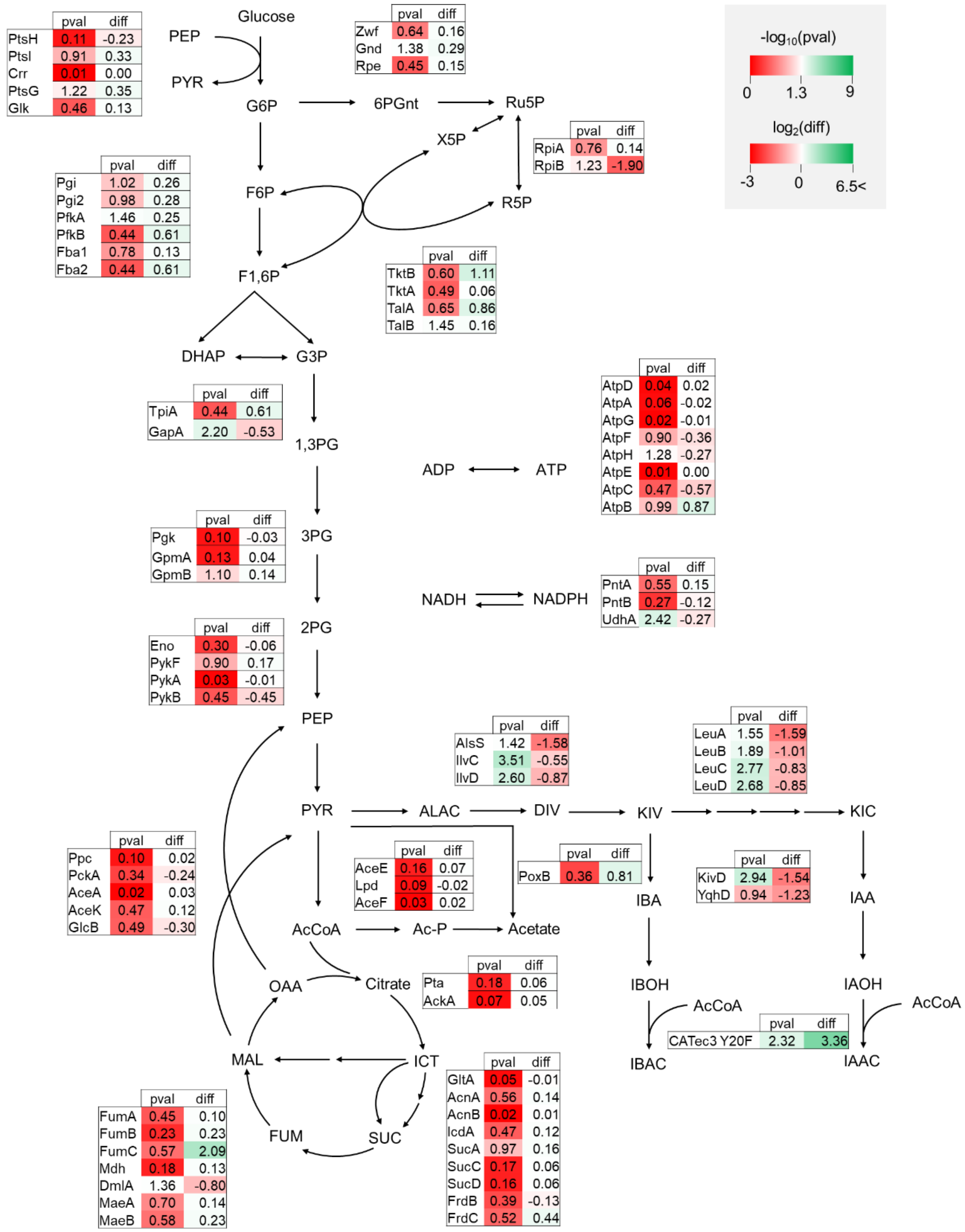

**Figure S3.** Isoamyl acetate titers of engineered strains with sequential gene deletions after 48 h culturing. Each data represents a mean ± 1 standard deviation from three biological replicates.

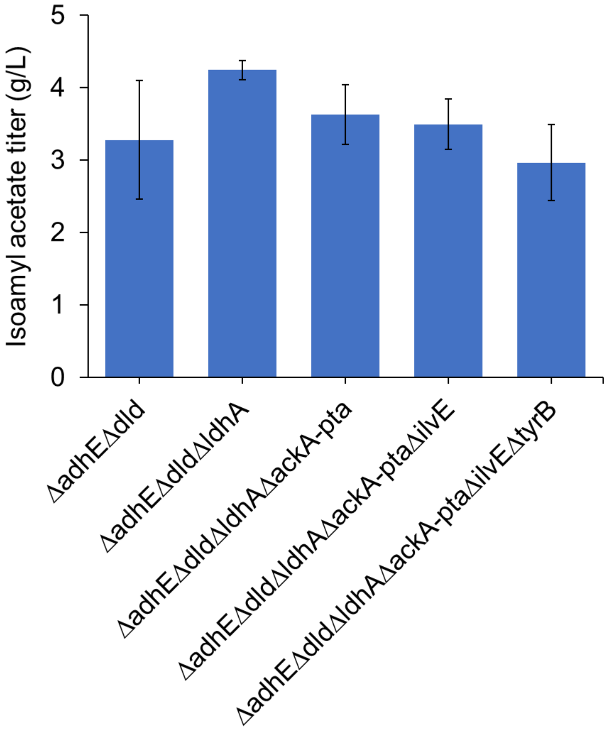

**Figure S4.** Growth of HSEC1311 under anaerobic and microaerobic conditions. HSEC1311 was cultured in M9 plus glucose medium with 0.1 mM IPTG. For the anaerobic condition, cell cultures were prepared in an anaerobic chamber and cultured in a 16 mL rubber sealed Balch tube. For the microaerobic condition, cells were cultured in a 125 mL screw capped flask. Each data represents a mean ± 1 standard deviation from three biological replicates.

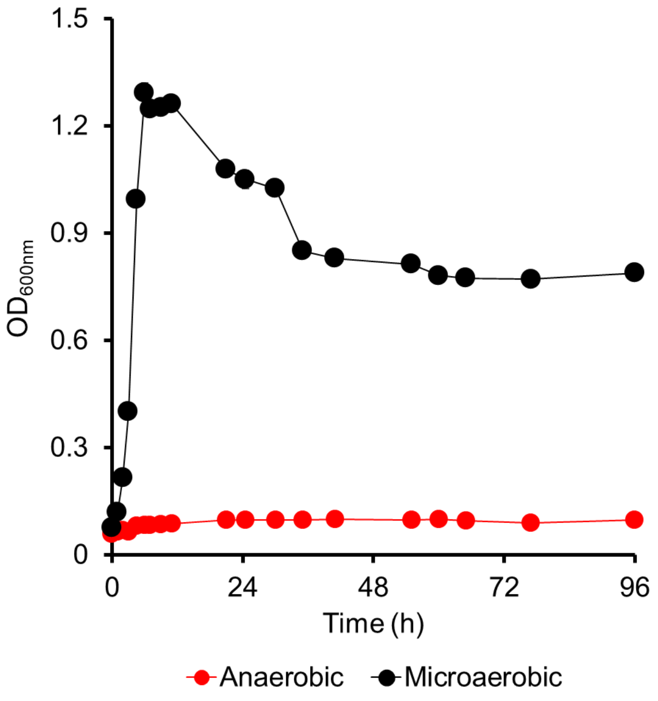
